# Multiple carotenoid-based signals are enhanced despite poor body condition in urban male and female Northern Cardinals (*Cardinalis cardinalis*)

**DOI:** 10.1101/2022.03.08.483494

**Authors:** Daniel T. Baldassarre, Haley S. Bedell, Kristie M. Drzewiecki, Brooke D. Goodman, Mya L. Mills, Denis A. Ramos

## Abstract

If humans aim to sustainably coexist with wildlife, we must understand how our activity impacts the communication systems of urban animal populations. We know much about the effects of anthropogenic noise on bird song, but relatively little about how avian visual signals are affected by urbanization. One way such an effect may occur if urbanization alters the food available to species with color based on carotenoids, which they must obtain from their diet. Over three years, we compared a comprehensive suite of visual signals in male and female Northern Cardinals (*Cardinalis cardinals*) in a rural and an urban population. We predicted that urban birds would have enhanced carotenoid-based signals as they likely have access to more carotenoids from invasive plants, especially honeysuckle (*Lonicera* spp.), that thrive in cities. We used reflectance spectrometry, digital image analyses, and avian visual models to quantify hue, saturation, and brightness of chest (male), underwing (female), and bill (male and female) signals. Compared to rural males, urban males had redder chest feathers in one year and redder bills in every year. Urban females had more saturated underwing color than rural females in every year. These color differences were sufficient to be distinguished by the avian visual system. Urbanization did not affect female bill color. Interestingly, urban birds had significantly reduced mass-related body condition compared to rural birds. These results show that both male and female urban birds can display enhanced carotenoid-based signals despite being in relatively poor condition. The consequences of this color enhancement are unknown, but it could affect the information content of the signals and the dynamics of the social and mating systems. These results stand in stark contrast to the predominant trend in birds of decreased color in urban areas and highlight the complex and varied potential effects of urbanization on animal communication.

---

Humans have drastically altered the natural landscape and it is imperative to understand how our activity affects animal populations if we aim to promote sustainable coexistence with wildlife (Seddon et al. 2016, Elmqvist et al. 2019). Beyond the negative impacts of human activities like direct exploitation (e.g., poaching, pet trade) and habitat destruction on wildlife population viability, humans can also more subtly alter evolutionary dynamics (Szulkin et al. 2020). For instance, considerable research demonstrates that humans can disrupt animal communication systems (Shannon et al. 2016, Slabbekoorn et al. 2018). In particular, urbanization has a myriad of impacts on signal production, transmission, and perception (Sepp et al. 2018, Sepp et al. 2020, Heinen-Kay et al. 2021). Urban environments present several challenges that may affect communication systems, including anthropogenic noise and light, heavy metal pollution, habitat fragmentation, and changes in resource availability (Heinen-Kay et al. 2021).

In birds, much of this work has focused on anthropogenic disruption of vocal communication, especially in urban areas (Slabbekoorn and den Boer-Visser 2006, Slabbekoorn 2013, Derryberry and Luther 2021). Less well understood is how visual signals may also be impacted by human development. Carotenoid-based color patches may be especially affected by urbanization as carotenoids cannot be synthesized de novo by birds and must be obtained from the diet (Hill 1991). One way urbanization can directly impact these signals, and hence the communication system, is if rural and urban areas differ in the access to, amount, or types of resources containing carotenoids. Moreover, carotenoids can undergo extensive physiological modification before being deposited in tissue (Goodwin 1984), leaving room for other more indirect effects of urbanization on color (e.g., via stress or parasites; Møller 2009, Hutton and McGraw 2016). Additionally, rural and urban habitats likely differ in lighting regime, which can also affect carotenoid-based color via selection on conspicuousness (Leveau 2021). Importantly, these mechanisms are not mutually exclusive and might either reduce or enhance color in urban populations. To date, research comparing carotenoid-based signals between rural and urban populations has found reduced ornamentation in urban populations of House Finches (*Haemorhous mexicanus*; Hasegawa et al. 2014, Giraudeau et al. 2015, Giraudeau et al. 2018) and Great Tits (*Parus major*; Grunst et al. 2020). However, urban birds could have more heavily-pigmented, enhanced carotenoid-based color if, for example, they have access to carotenoid-rich invasive plants that thrive in disturbed landscapes (Hobbs 2000, Vilà and Ibáñez 2011). In the midwestern and northeastern United States, invasive honeysuckle (*Lonicera* spp.) has been implicated in the enhancement of carotenoid-based signals in a variety of bird species (e.g., Northern Cardinal, *Cardinalis cardinalis*; Cedar Waxwing, *Bombycilla cedrorum*; Baltimore Oriole, *Icterus galbula*; Northern Flicker, *Colaptes auratus*; Mulvihill et al. 1992, Hudon et al. 2013, Hudon et al. 2017, Hudon and Mulvihill 2017). *Lonicera* thrives in urban areas (Borgmann and Rodewald 2005, Rodewald 2005, Leston and Rodewald 2006), providing a potential link between urbanization and enhanced color. However, to our knowledge, no study has found enhanced carotenoid-based signals in an urban population of any species. Quantifying the color of rural and urban populations of a species that occurs in both habitats provides a powerful framework to test this idea.

The Northern Cardinal (hereafter “cardinal”) is well suited to such a study because it readily occurs in both rural and urban areas, and both males and females exhibit carotenoid-based color patches that are displayed during various territorial, courtship, and pre-copulatory interactions and likely function as signals (Linville et al. 1998, Wolfenbarger 1999a, Wolfenbarger 1999b, Jawor et al. 2003, Jawor et al. 2004, Jawor and Breitwisch 2004). Males have bright red plumage overall and orange-red bills, while females have pink-red underwing patches (hereafter “wing”) and orange-red bills. The putative functions of these colors as signals are complex and variable across studies. For example, males with redder chest color had higher quality territories (Wolfenbarger 1999a, Rodewald et al. 2011) and reproductive success (Wolfenbarger 1999a), but females did not prefer redder males (Wolfenbarger 1999b). Males and females mated assortatively based on chest, wing, and bill color (Jawor et al. 2003); and female wing and bill color co-varied with body condition (Jawor et al. 2004). The color of carotenoid-based plumage color in males is related to carotenoid availability in fruits, which they favor during molt in late summer and fall (Linville and Breitwisch 1997); and captive studies confirm that carotenoid intake affects plumage color (McGraw and Hill 2001). Cardinals have been studied in an urban context where males, but not females, had reduced brightness as urbanization increased across a gradient in Ohio (Jones et al. 2010), which was attributed to the generally negative effects of urban development. In Florida, rural and urban males did not differ in plumage color (Leigh 2012). Unlike plumage, the effects of urbanization on bill color have not been studied in this or any species of which we are aware.

Our goal was to compare multiple carotenoid-based signals (chest, bill, and wing color) in rural and urban males and females while accounting for the sensitivity of the avian visual system, which has not been implemented in this species. We hypothesized that urbanization would affect the color of these signals, specifically that urban birds would have enhanced signals (e.g., redder hue, increased saturation) compared to rural birds, potentially due to increased access to carotenoids. We also explored relationships between mass-related body condition, urbanization, and color.

## Methods

### Field sites and data collection

We studied cardinals in 2 populations in Central New York that we chose because they differed drastically in the extent of human development (Fig. 1). The rural site, Rice Creek Field Station (43.43°N, 76.55°W; Fig. 1a), was located outside of Oswego (population 17,470), while the urban field site, Barry Park (43.03°N, 76.12°W; Fig. 1b), was located within the city limits of Syracuse (population 142,874). The 2 sites were separated by 57 km. At each site, we captured wild individuals using a territory-based targeted mist net approach between May and July in 2019, 2020, and 2021. We did not re-capture any individuals across years. Although we did not know the precise age of most individuals, we limited our analyses to those we could age as “after-hatching-year” (i.e., at least one-year-old) based on plumage. No captured individuals had yet initiated their prebasic molt.

**Figure 1.**
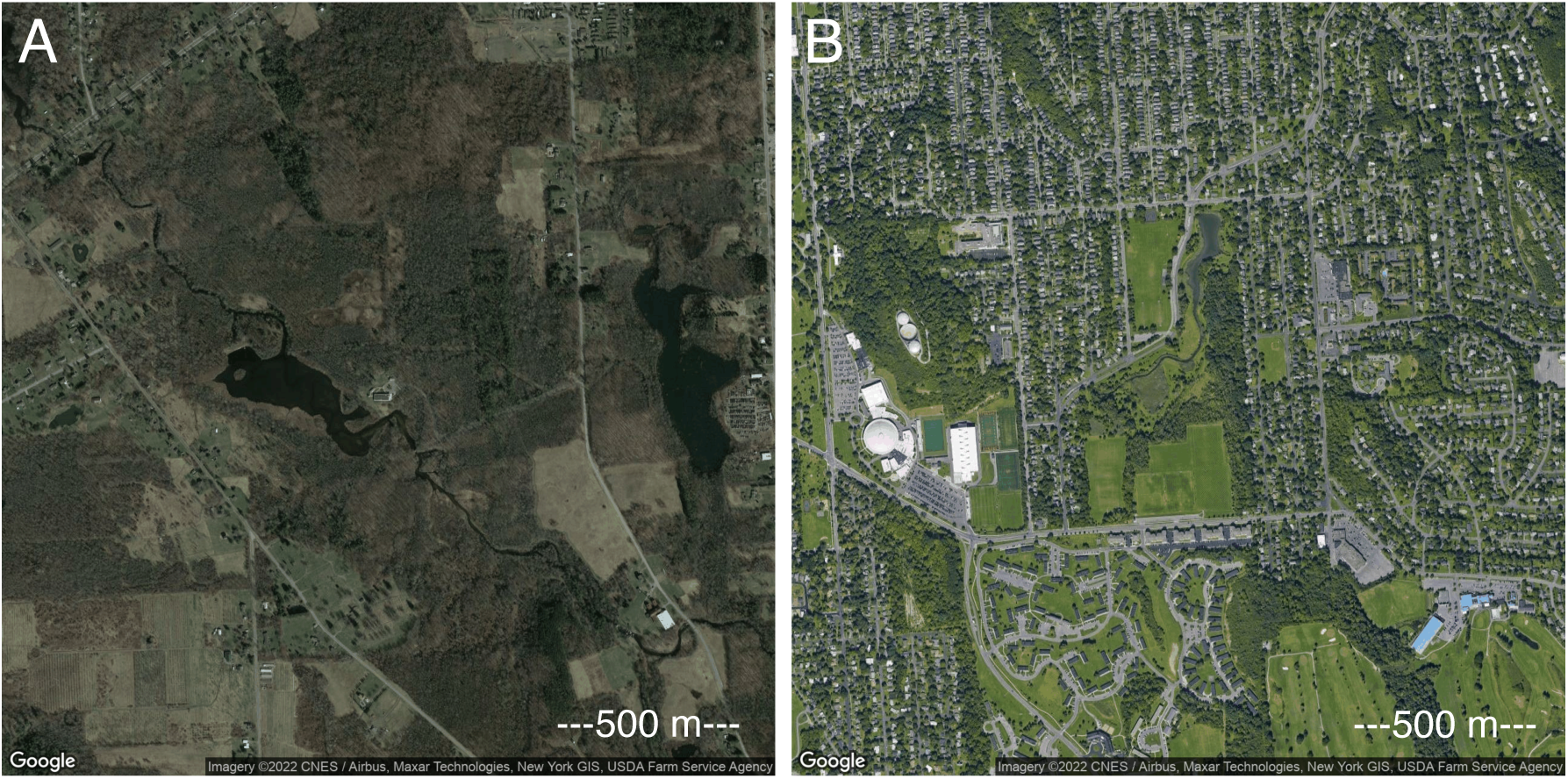
A map of the rural (Rice Creek Field Station, a) and urban (Barry Park, b) field sites.

To quantify body condition, we measured mass to the nearest 0.5 g with a spring scale, and tarsus length to the nearest 0.01 mm with digital calipers. Both populations were in the midst of breeding, but we did not have precise information on nest stage for most individuals. For males and females separately, we regressed mass on tarsus length. Males exhibited a significant positive relationship between tarsus length and mass (*F*_1,63_ = 18.0, *P* < 0.001), so we extracted model residuals and used them as an index of body condition. This index described whether an individual was heavier (positive values) or lighter (negative values) than would be predicted by body size. There was no significant relationship between female mass and tarsus length (*F*_1,23_ = 0.7, *P* = 0.41), so we used mass as an index of body condition.

We collected 10 feathers from the same central region of the chest of each captured male for analysis via reflectance spectrometry (rural *n* = 43, urban *n* = 22). In the lab, we mounted the feathers in an overlapping pattern on black construction paper (Strathmore Artagain Coal Black) and measured ultraviolet (UV) and visible light reflectance from 300-700 nm using an Ocean Optics Jaz reflectance spectrometer and a PX2 lamp. The probe was mounted at a 90° angle in a block that held the probe tip 5 mm from the feather surface and excluded ambient light. Between each sample, we calibrated the spectrometer using a WS-1 white standard. We took 3 measurements across different regions of the feather sample and averaged the reflectance curves before analysis (Supplemental Fig. S1). To quantify male bill (rural *n* = 40, urban *n* = 21), female bill (rural *n* = 16, urban *n* = 9), and female wing (rural *n* = 15, urban *n* = 9) color, we took photographs in the field using a Canon EOS Rebel T6 digital camera (Supplemental Fig. S2). Variation in sample size across signals was due to some photographs being unusable. We used automatic settings because there were no systematic differences in ISO (*t* = -0.7, df = 47.3, *P* = 0.5) or aperture (*t* = -0.8, df = 42.9, *P* = 0.41) between sites. Each photograph contained color standards and a scale bar (X-Rite ColorChecker Passport) and was taken under diffuse natural sunlight without shadows. The inclusion of color standards in each photo allowed us to compare among images controlling for variation in ambient light. We took 3 photos of each color patch that varied in exposure (one stop under-exposed, normal, and one stop over-exposed) and then selected the best-exposed, sharpest image for analysis.

### Color analyses

We analyzed reflectance curves of male chest feathers using the program *pavo 2* (Maia et al. 2019) in R 3.6.0 (R Core Team 2019). We obtained quantum catch values for each of the 4 avian single cones using the Blue Tit (*Cyanistes caeruleus*) visual model (plus double-cone sensitivity for luminance, hereafter “brightness”) and idealized illuminant D65. We then used the quantum catch values to represent each color as a point in tetrahedral color space where the vertices represented stimulation of the u, s, m, and l cones. From the color space, we extracted the angle of the color vector relative to the x-y (theta) and z (phi) plane, which together describe hue. Theta and phi were highly correlated (*R* = -0.94, *P* < 0.001), so we only used theta for subsequent analyses. To quantify saturation, we used the distance between the point and the achromatic center relative to the maximum possible distance. Together these measurements described hue, saturation, and brightness of the chest feathers accounting for UV reflectance and the sensitivity of the avian visual system.

We analyzed digital photographs of male and female bills and female wing patches using the Quantitative Color Pattern Analysis (QCPA; van den Berg et al. 2020) framework in the ImageJ 1.53a (Schneider et al. 2012) plugin micaToolbox (Troscianko and Stevens 2015). We first generated a cone catch model accounting for the camera’s photosensitivity, the Blue Tit visual model, and idealized illuminant D65. This calibration was achieved by measuring an image of the color standards, which have known reflectance values. The camera was not UV-sensitive, so these analyses were restricted to the visible spectrum (400-700 nm). However, these orange-red color patches reflect predominantly long, visible wavelengths, where most of the variation occurs (Supplemental Fig. S1). Once a cone catch model was built, we converted each image into a normalized multispectral stack standardized by the 3% and 92% reflectance standards and selected regions of interest to measure (Supplemental Fig. S2a–b). We then ran each image through the cone catch model, followed by the QCPA framework that applies spatial acuity and viewing distance correction and then uses the Receptor Noise Limited (RNL) model (Vorobyev and Osorio 1998) to reduce the image into discernable color clusters that can be measured (Supplemental Fig. S2c–d). We used Gaussian acuity correction with 6 cycles per degree and a simulated viewing distance of 0.5 m. We used RNL ranked filtering with the following settings: iterations: 5, radius: 5, falloff: 3. We used RNL clustering with the following settings: visual system Weber fractions: Blue Tit 0.05, luminance Weber fraction: 0.1, color JND threshold: 2, luminance JND threshold: 3, loops: 20, radius multiplier: 2, minimum cluster size: 10. For each color cluster produced by QCPA, we extracted the quantum catch values for the 3 avian single cones (excluding the UV-sensitive cone) and the double cone (i.e., brightness). When an image was split into more than one cluster by QCPA, we computed a weighted average of each quantum catch value accounting for the area of each cluster. Using *pavo 2*, we converted these values into a color point in triangular color space where the vertices represented stimulation of the s, m, and l cones. From the color space, we extracted the single angle of the color vector (theta), which describes hue. To quantify saturation, we used the distance between the point and the achromatic center. Together these measurements described hue, saturation, and brightness of the male and female bill and female wing patches considering the visible spectrum and the sensitivity of the avian visual system. Note that although both the tetrahedral and triangular color spaces produce angle theta to describe hue, the difference in geometry is such that higher values for male chest correspond to more orange hue, while they correspond to redder hue for male and female bill and female wing.

If we discovered a significant difference in color variables between factors, we further explored whether these differences would be recognizable by the signal receiver. We input the quantum catch values into the noise-weighted color distance (dS) calculator in *pavo 2* using the following settings: Weber fraction: 0.05, noise: neural, photoreceptor densities: u = 1, s = 2, m = 2, l = 4. We compared the resulting values to dS = 1, or the perceptual threshold indicating colors that can be distinguished by the avian visual system (Vorobyev and Osorio 1998).

### Statistical analyses

Because hue, saturation, and brightness represent different and potentially informative aspects of a color, we analyzed them as separate response variables for each signal. We considered site (rural vs. urban) and year (2019, 2020, 2021) as factors because color could vary depending on habitat characteristics or could change systematically due to environmental differences over time. When checking for collinearity among predictors, we found a significant correlation between site and body condition using a linear model (see Results), and thus did not include body condition as a factor in these models. We used a model selection approach, with all possible combinations of factors as predictors in a linear model, as well as an intercept-only model. We ranked models by Akaike’s Information Criterion corrected for small sample sizes (AICc) and considered the model with the lowest AICc score to be best supported by the data. We considered models with ΔAICc < 2 to have similar fits compared to the top model. Additionally, we used log likelihoods and normalized Akaike weights to compare models. We conducted post-hoc comparisons using Tukey’s HSD corrected for multiple comparisons. To determine whether dS values between factors were larger than the avian perceptual threshold, we tested whether the average pairwise dS value of all cross-factor comparisons (e.g., between every rural and urban color) was significantly greater than one using a one-sample *t*-test. Analyses were performed using R 3.6.0 (R Core Team 2019) and the *MuMIn* package (Bartón 2020).

## Results

The top model for male chest hue included an interaction between site and year (Supplemental Table S1). There was a significant effect of year (*F*_2,59_ = 18.7, *P* < 0.001) and the interaction between site and year (*F*_2,59_ = 5.0, *P* = 0.01). Male chest hue in 2020 was higher (more orange) than 2019 and 2021 (both *P* < 0.001), and urban hue was lower (redder) than rural hue in 2020 (*P* = 0.03, Fig. 2a). The top model for male chest saturation included year only (Supplemental Table S1). There was a significant effect of year (*F*_2,62_ = 8.7, *P* < 0.001) such that saturation in 2019 was higher than 2020 (*P* = 0.003) and 2021 (*P* = 0.01, Fig. 2b). Similarly, the top model for male chest brightness also included year only (Supplemental Table S1). There was a significant effect of year (*F*_2,62_ = 13.4, *P* < 0.001) such that brightness in 2021 was lower than 2019 and 2020 (both *P* < 0.001, Fig. 2c). In 2020, the significant difference in male chest hue between the rural and urban site resulted in an average pairwise dS value that was significantly greater than the perceptual threshold (*t* = 6.0, df = 35, *P* < 0.001, Fig. 4a).

**Figure 2.**
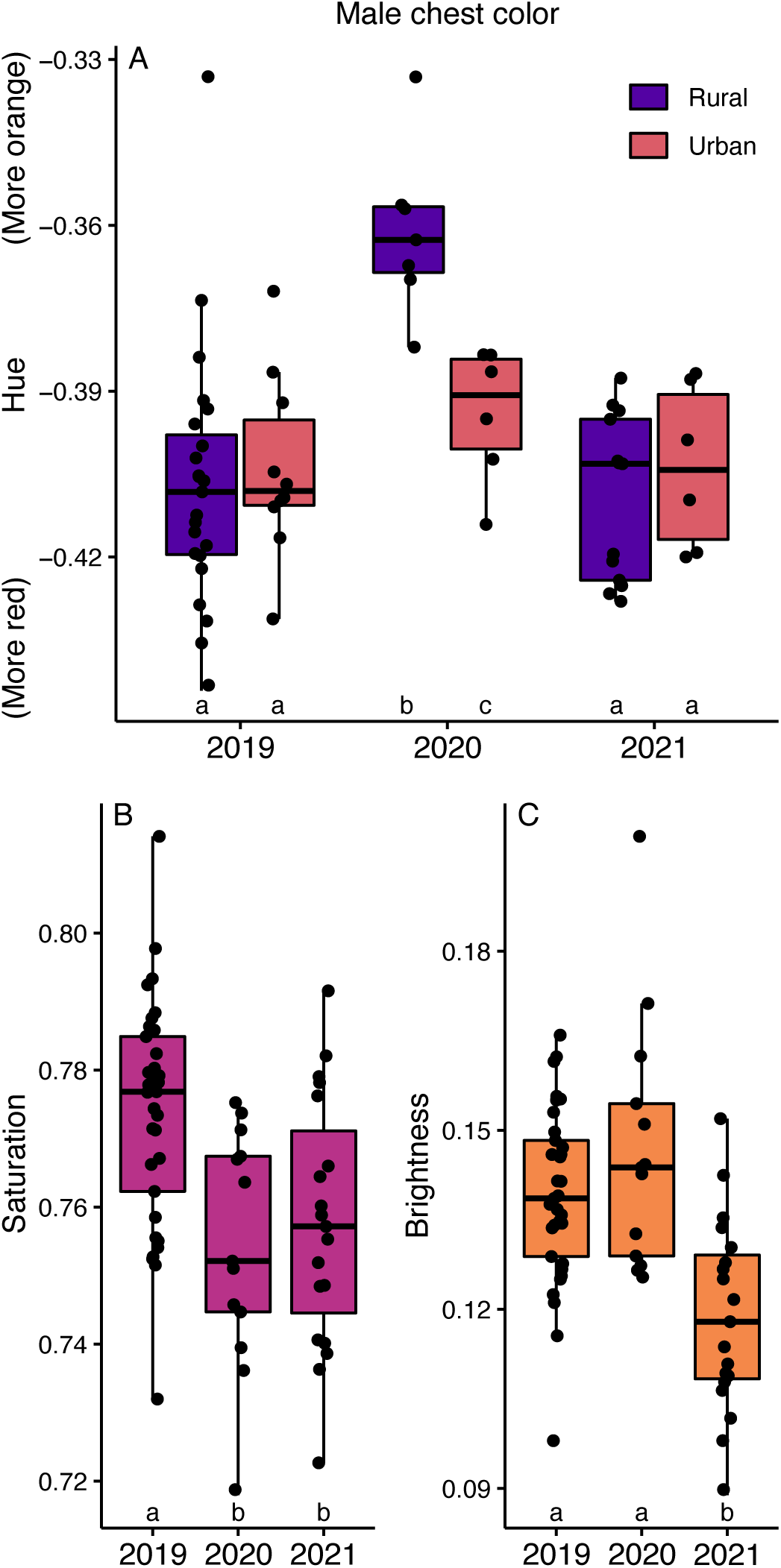
Variation in male chest hue (a), saturation (b), and brightness (c) as a function of urbanization and/or year. There was no significant effect of site on saturation or brightness, so only the effect of year is shown. Lowercase letters indicate which factors are (different letter) or are not (same letter) significantly different from each other.

The top model for male bill hue included site only (Supplemental Table S2). There was a significant effect of site (*F*_1,59_ = 12.4, *P* = 0.001) such that urban hue was higher (redder) than rural hue (Fig. 3a). There were no significant effects of any factors on male bill saturation or brightness, as the top models were indistinguishable from an intercept-only model (Supplemental Table S2). The significant difference in male bill hue between the rural and urban site resulted in an average pairwise dS value that was significantly greater than the perceptual threshold (*t* = 35.5, df = 839, *P* < 0.001, Fig. 4a). There were no significant effects of any factors on female bill hue, saturation, or brightness (Supplemental Table S3). For female wing saturation, the top model included site only (Supplemental Table S4). There was a significant effect of site (*F*_1,22_ = 4.9, *P* = 0.04) such that urban saturation was higher than rural saturation (Fig. 3b). There were no significant effects of any factors on female wing hue or brightness (Supplemental Table S4). The significant difference in female wing saturation between the rural and urban site resulted in an average pairwise dS value that was significantly greater than the perceptual threshold (*t* = 11.1, df = 134, *P* < 0.001, Fig. 4a).

**Figure 3.**
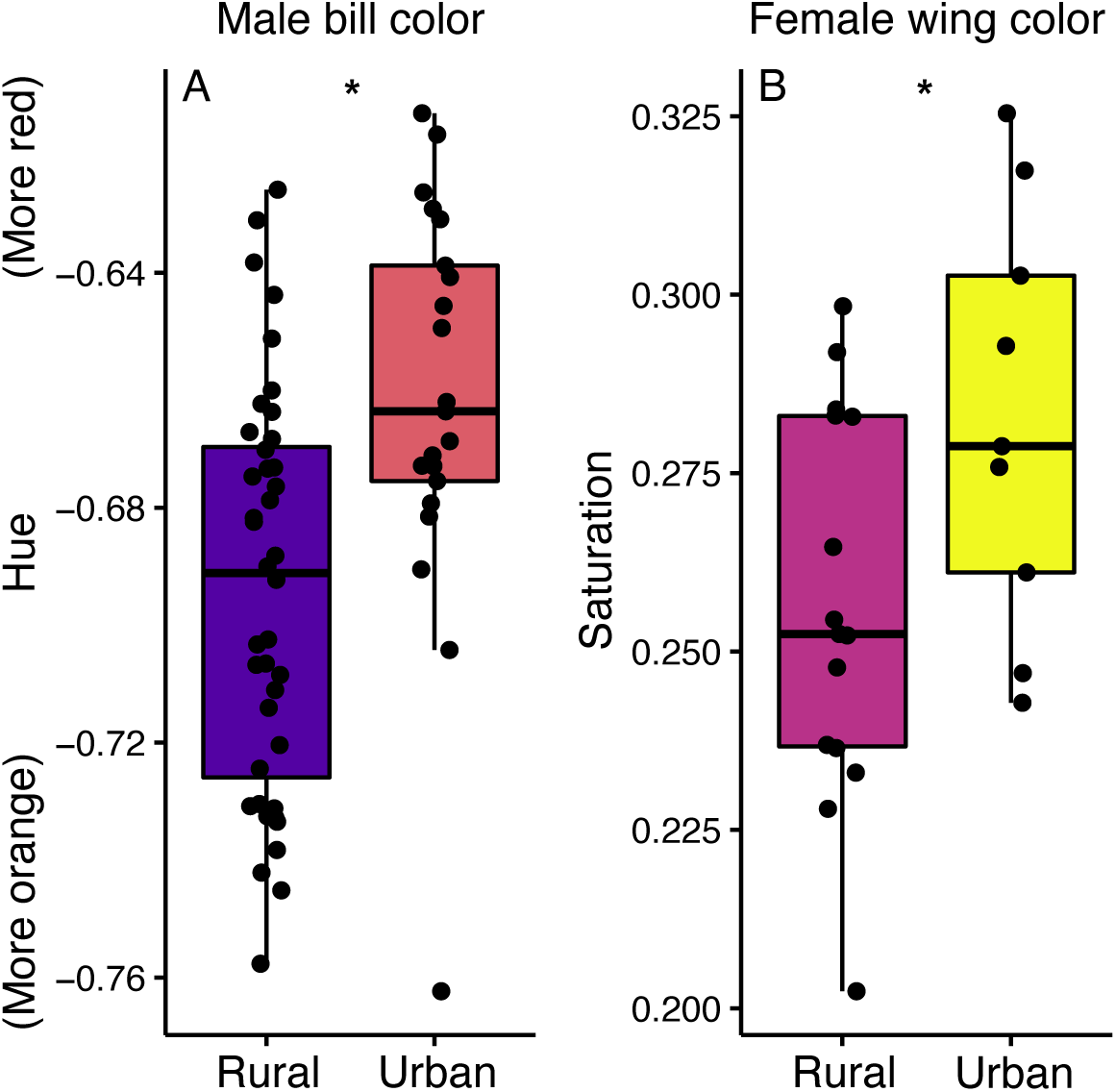
Variation in male bill hue (a) and female wing saturation (b) as a function of urbanization. Asterisks separate factors that are significantly different.

**Figure 4:**
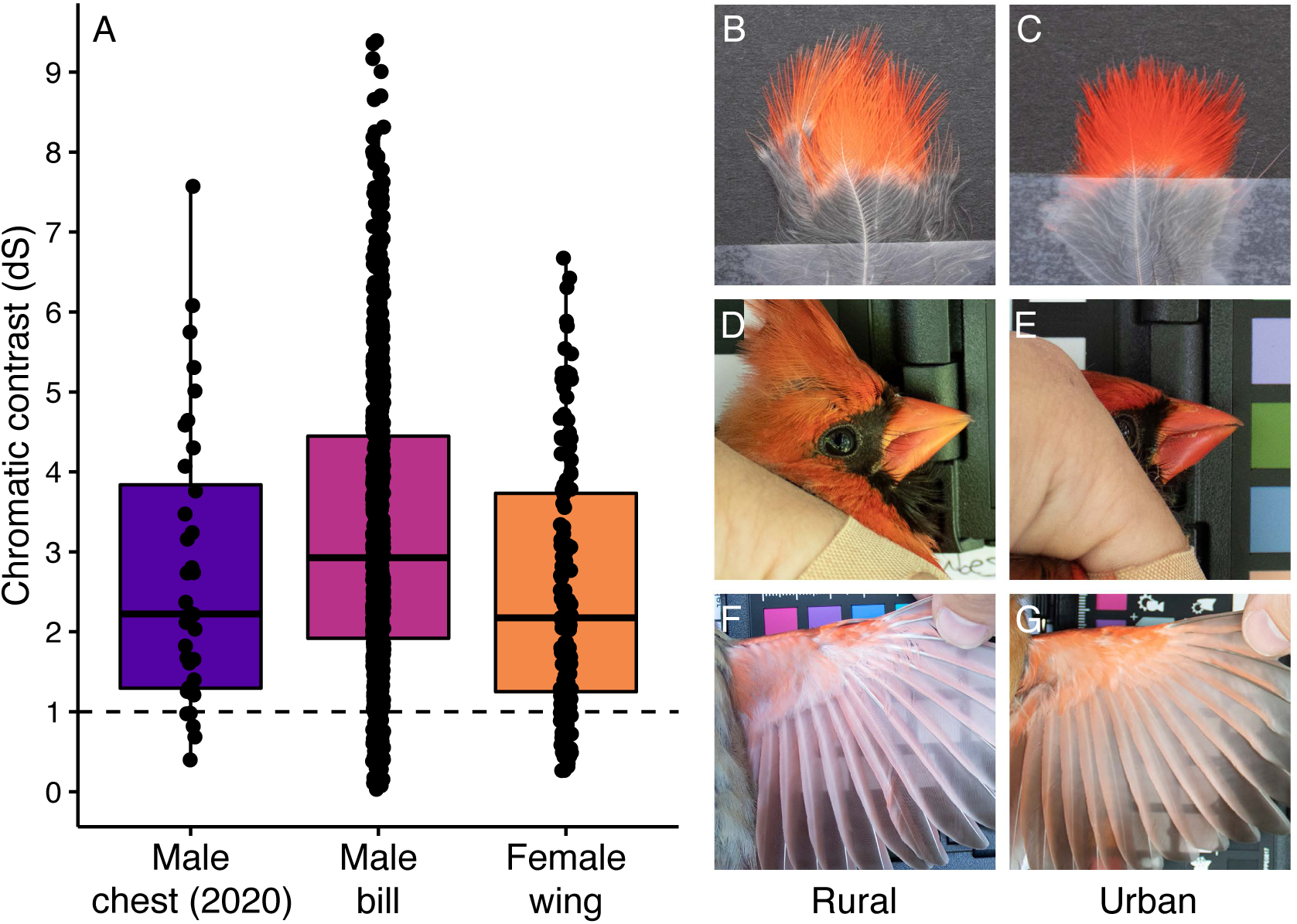
Pairwise chromatic contrast between rural and urban male chest (2020 only), male bill, and female wing (a). The dashed line indicates dS = 1, or the perceptual threshold above which colors can be distinguished by the avian visual system. Photos exemplifying the average rural and urban male chest (b, c), male bill (d, e), and female wing (f, g) signals.

Mass-related body condition of both males and females was affected by site. Male body condition score was higher in rural compared to urban birds (*F*_1,63_ = 14.9, *P* < 0.001, Figure 5a), and female mass was higher in rural compared to urban birds (*F*_1,23_ = 4.5, *P* = 0.05, Figure 5b).

**Figure 5.**
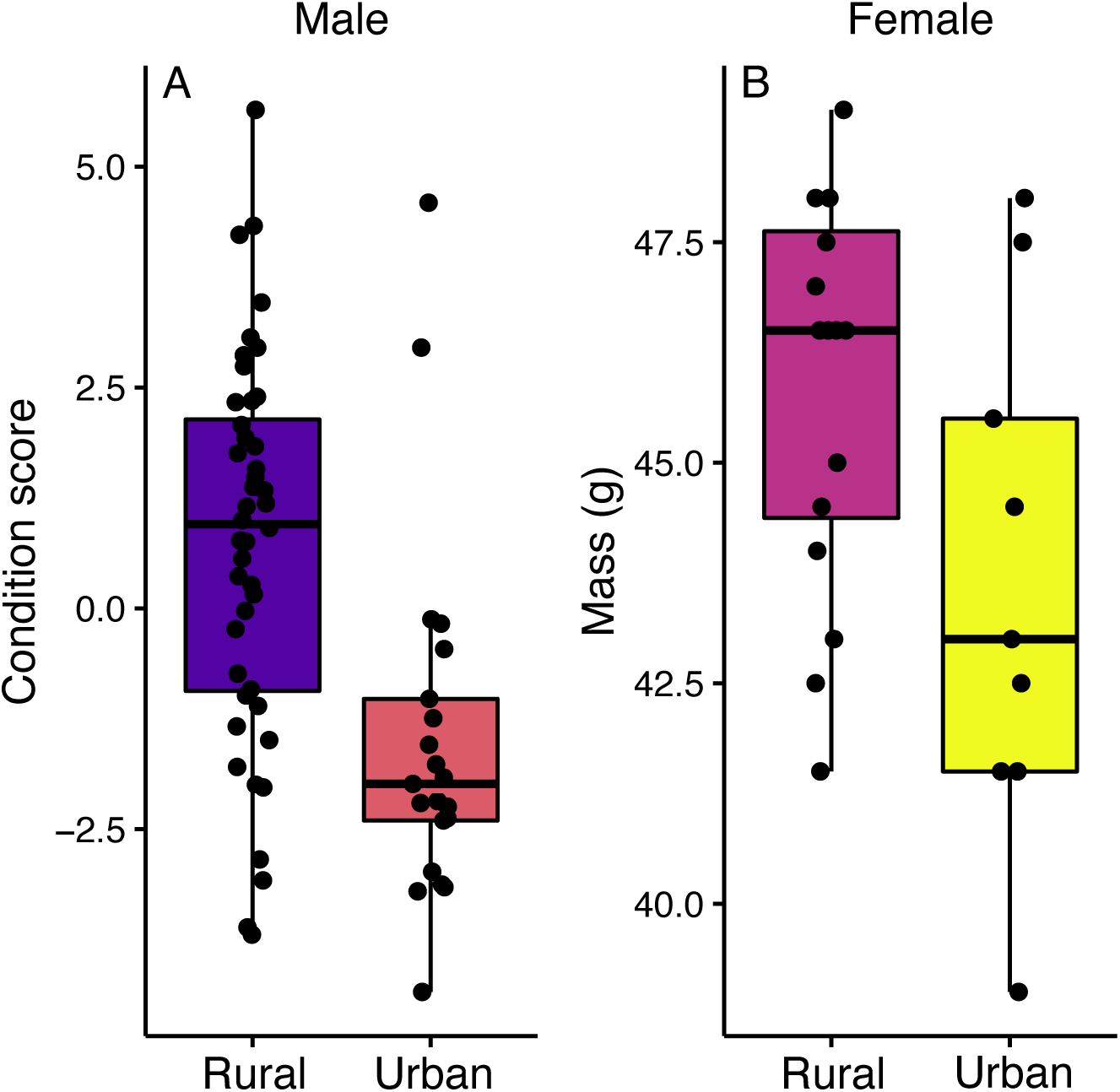
Variation in male body condition score (a) and female mass (b) as a function of urbanization.

## Discussion

The color of multiple carotenoid-based signals was enhanced in males and females compared to rural birds. Specifically, these colors were redder or more saturated in urban birds (Fig. 2–3), despite these individuals being in relatively poor mass-related body condition compared to rural birds (Fig. 5). Furthermore, although subtle to the human eye, these color differences were large enough to be distinguished by the avian visual system (Fig. 4). These findings suggest that urban living may reduce mass-related body condition, but that urban birds have enough access to carotenoids that they are nonetheless able to produce what are typically thought of as costly honest signals. In rural habitats, carotenoids may be a much more limiting resource, such that rural birds have reduced ornamentation overall, but color may be a more honest signal of condition. Preliminary analyses have found weak or no relationships between body condition and ornament color within either population, but we are hesitant to over-interpret these results due to low sample sizes. Our findings are in line with previous work on cardinals that found a dampened relationship between body condition, plumage color, and reproductive success as urbanization increased (Jones et al. 2010, Rodewald et al. 2011). Whether cardinals attend to the color variation documented here, whether that variation is linked to reproductive success, and whether the urban color enhancement disrupts the typical signaling dynamics are unknown in our populations. However, our study stands apart in that we found a distinct enhancement of multiple carotenoid-based signals in urban birds of both sexes, apparently not associated with increased mass-related body condition. This is the first such result we are aware of, as the overwhelming trend in the literature is for reduced ornamentation in urban areas, even among melanin-based or structural colors (Sepp et al. 2018, Sepp et al. 2020, Heinen-Kay et al. 2021).

A closer inspection of the chest color results supports the idea that increased and/or consistent access to carotenoids in urban areas could be what promoted the color enhancement. In 2020 when we found a significant difference in chest hue between rural and urban males, overall hue was more orange than in other years, but this difference was greater in the rural habitat, which drove the rural-urban difference (Fig. 2A). This pattern suggests that carotenoid availability may fluctuate more widely over time in the rural habitat, but that urban birds have a more stable carotenoid supply that allows them to maintain redder plumage. Leigh (2012) made a similar interpretation based on urban male cardinals in Florida exhibiting less among-individual variation in chest brightness than rural males. We suggest that the relative abundance of exotic fruiting shrubs, especially *Lonicera* spp., in urban areas (Borgmann and Rodewald 2005, Borgmann and Rodewald 2005, Leston and Rodewald 2006) is responsible for the pattern we observed. Although we have not quantified plant diversity at our sites, anecdotally, *Lonicera* is much more established at the urban site compared to a more diverse vegetation community at the rural site (DTB, 2019, pers. observ.). We did not choose these sites a priori based on this difference. Furthermore, an increase in sampling sites would help determine if the observed color variation is due to *Lonicera* abundance specifically or other potential differences between rural and urban sites. Additionally, work is ongoing to quantify carotenoids in blood plasma and feathers to determine more conclusively if urban birds ingest and/or deposit more carotenoids than rural birds.

Whether or not carotenoids are more available to urban birds, it is important to note that many other unmeasured factors may influence whether they are acquired and deposited in tissue. For example, even if carotenoid-based color is a condition-dependent trait in cardinals, our mass-related measures of body condition offer only a small glimpse into what might be physiologically relevant to true condition. Rural and urban birds may differ in other more meaningful measures of condition such as immune function or stress. Wright and Fokidis (2016) found that urban male cardinals had lower circulating corticosterone and a lower stress response than rural males. Urban habitats may be less stressful for cardinals and/or they may modulate their stress response such that they can more efficiently metabolize and deposit carotenoids in their tissues regardless of availability (Giraudeau et al. 2015). Related to this idea, in a suite of Western Palearctic species, Møller (2009) found that urban birds had a stronger immune response than rural birds, which might free up carotenoids to be deposited in tissue. Additionally, if the lighting conditions and/or visual background differs between sites, urban and rural birds may experience disruptive selection to maximize the conspicuousness of their visual signals (Dalrymple et al. 2018, Leveau 2019). In summary, it is difficult with the present data to determine the mechanism of the observed color variation, and there are several, non-mutually exclusive possibilities.

It is unclear why urban birds were in worse mass-related body condition than rural birds. Potential explanations include limited access to protein-rich insect prey, benefits of reduced mass for predator evasion, and adaptive changes in nestling development. In cardinals, previous studies have found little to no effect of urbanization on similar body condition metrics (Rodewald and Shustack 2008, Jones et al. 2010) or even that urban birds were in better condition than rural birds (Wright and Fokidis 2016). However, the potential impacts of urbanization on body condition may be site-specific, and there is evidence in other species of a negative effect (Jiménez-Peñuela et al. 2019). Additionally, urban birds may not need to accumulate energy reserves in the form of body mass in urban areas where food availability is more predictable and nutritional stress is not as severe as in rural areas (Salleh Hudin et al. 2016). Regardless of the cause, we found that despite this reduction in mass-related body condition, urban birds were still able to produce enhanced color signals compared to rural birds.

We found an enhancement of both feather and bill color in urban birds. The timing of production of these two signal types has different implications for the relationship with carotenoid availability. Cardinals molt once per year after the breeding season, so chest and wing color should mainly be influenced by carotenoid deposition at that time only. However, it is possible that the color variation is due to differences between rural and urban birds in the extent to which feather color wears over time (e.g., due to differences in sun exposure). In contrast to plumage, carotenoid-based bill color can vary much more dynamically and over shorter time periods (Rosen and Tarvin 2006, Ardia et al. 2010). If the rural-urban color difference in bill color is driven by carotenoid availability, these results may suggest that urban birds have greater access to carotenoids not only during molt, but potentially throughout the breeding season (when these birds were captured and photographed). Since *Lonicera* fruits are mainly available in late summer and fall, carotenoids producing bill color may come from different sources (e.g., insect prey), although fruits can ripen as early as June (DTB, 2020, pers. observ.). However, the extent to which cardinal bill color changes over time is unknown, and color measured during the summer breeding season may well have resulted from carotenoids ingested and sequestered previously. Importantly, the factors influencing color (e.g., carotenoid availability, body condition, stress) may vary across the signals measured in this study.

## Conclusion

Urbanization typically dulls or reduces the extent of color in birds (Sepp et al. 2018, Leveau 2019, Heinen-Kay et al. 2021). Here we show instead that both male and female urban cardinals had significantly enhanced color of multiple carotenoid-based signals compared to rural birds. We suggest this enhancement is potentially due to the widespread and stable availability of carotenoids in exotic plants that thrive in disturbed areas, although there are other possible explanations, and these mechanisms deserve additional investigation. If carotenoids are a limiting resource in rural but not urban habitats, the reliability of these signals as honest indicators of condition may deteriorate (Jones et al. 2010, Rodewald et al. 2011). In support of this idea, we also found that urban birds were in worse mass-related body condition compared to rural birds, suggesting a decoupling of condition and color. Together, these results underscore the complex and subtle impacts of urbanization on wildlife. Not only can urbanization negatively affect populations by destroying and fragmenting habitat, it may also disrupt communication systems and potentially alter the evolutionary dynamics of signals.

## Acknowledgments

Susan DeVries, Sylvia Halkin, Jodie Jawor, Gavin Leighton, and Daniel Shustack provided helpful comments on early drafts of this manuscript. Comments from anonymous reviewers greatly improved the manuscript. Shyla Luna helped collect data in the field. We are grateful to Syracuse University and the Onondaga County Department of Water Environment Protection for access to Barry Park, and to Alan Harris for logistical support at Rice Creek Field Station. Lauren Silvernail provided childcare while fieldwork was being conducted. This research was supported by Rice Creek Associates and the Scholarly and Creative Activities Committee at SUNY Oswego. This research was approved by the SUNY Oswego Institutional Animal Care and Use Committee (protocol 2018.05) and conducted under appropriate federal (banding permit 24047), state (NYS DEC License 229), and local permits.

