## Supplemental Material for "Multiple carotenoid-based signals are enhanced despite poor body condition in urban male and female Northern Cardinals (*Cardinalis cardinalis*)"

**Table S1.** Summary of model selection results examining male chest hue, saturation, and brightness ($n$ = 65). Only models with ΔAICc < 5 are shown.

| Response | Factors | $k$ | logLik | ΔAICc | Weight |
|---|---|---|---|---|---|
| hue | site*year | 7 | 170.22 | 0 | 0.86 |
| hue | year | 4 | 164.76 | 3.63 | 0.14 |
| saturation | year | 4 | 173.58 | 0 | 0.66 |
| saturation | site+year | 5 | 173.58 | 2.35 | 0.21 |
| saturation | site*year | 7 | 175.62 | 3.23 | 0.13 |
| brightness | year | 4 | 175.87 | 0 | 0.58 |
| brightness | site*year | 7 | 178.50 | 2.05 | 0.21 |
| brightness | site+year | 5 | 176.01 | 2.07 | 0.21 |

**Table S2.** Summary of model selection results examining male bill hue, saturation, and brightness ($n$ = 61). Only models with ΔAICc < 5 are shown.

| Response | Factors | $k$ | logLik | ΔAICc | Weight |
|---|---|---|---|---|---|
| hue | site | 3 | 120.41 | 0 | 0.50 |
| hue | site+year | 5 | 122.53 | 0.43 | 0.40 |
| hue | site*year | 7 | 123.61 | 3.29 | 0.10 |
| saturation | site | 3 | 115.65 | 0 | 0.29 |
| saturation | intercept | 2 | 114.51 | 0.07 | 0.28 |
| saturation | site+year | 5 | 117.69 | 0.58 | 0.22 |
| saturation | year | 4 | 116.48 | 0.63 | 0.21 |
| brightness | intercept | 2 | 116.29 | 0 | 0.39 |
| brightness | year | 4 | 118.24 | 0.62 | 0.29 |
| brightness | site | 3 | 116.68 | 1.44 | 0.19 |
| brightness | site+year | 5 | 118.62 | 2.24 | 0.13 |

**Table S3.** Summary of model selection results examining female bill hue, saturation, and brightness ($n$ = 25). Only models with ΔAICc < 5 are shown.

| Response | Factors | $k$ | logLik | ΔAICc | Weight |
|---|---|---|---|---|---|
| hue | intercept | 2 | 47.09 | 0 | 0.50 |
| hue | year | 4 | 49.10 | 1.43 | 0.25 |
| hue | site | 3 | 47.35 | 2.07 | 0.18 |
| hue | site+year | 5 | 49.41 | 3.98 | 0.07 |
| saturation | year | 4 | 50.13 | 0 | 0.43 |
| saturation | intercept | 2 | 47.29 | 0.23 | 0.38 |
| saturation | site | 3 | 47.30 | 2.81 | 0.10 |
| saturation | site+year | 5 | 50.15 | 3.12 | 0.09 |
| brightness | year | 4 | 46.56 | 0 | 0.45 |
| brightness | intercept | 2 | 43.60 | 0.47 | 0.36 |
| brightness | site | 3 | 43.60 | 3.07 | 0.10 |
| brightness | site+year | 5 | 46.56 | 3.15 | 0.09 |

**Table S4.** Summary of model selection results examining female wing hue, saturation, and brightness ($n$ = 24). Only models with ΔAICc < 5 are shown.

| Response | Factors | $k$ | logLik | ΔAICc | Weight |
|---|---|---|---|---|---|
| hue | intercept | 2 | 42.30 | 0 | 0.51 |
| hue | site | 3 | 42.93 | 1.37 | 0.26 |
| hue | year | 4 | 43.97 | 2.21 | 0.17 |
| hue | site+year | 5 | 44.57 | 4.22 | 0.06 |
| saturation | site | 3 | 52.73 | 0 | 0.75 |
| saturation | intercept | 2 | 50.34 | 2.15 | 0.25 |
| brightness | year | 4 | 27.03 | 0 | 0.33 |
| brightness | intercept | 2 | 24.21 | 0.11 | 0.31 |
| brightness | site+year | 5 | 28.10 | 1.09 | 0.19 |
| brightness | site | 3 | 24.94 | 1.27 | 0.17 |

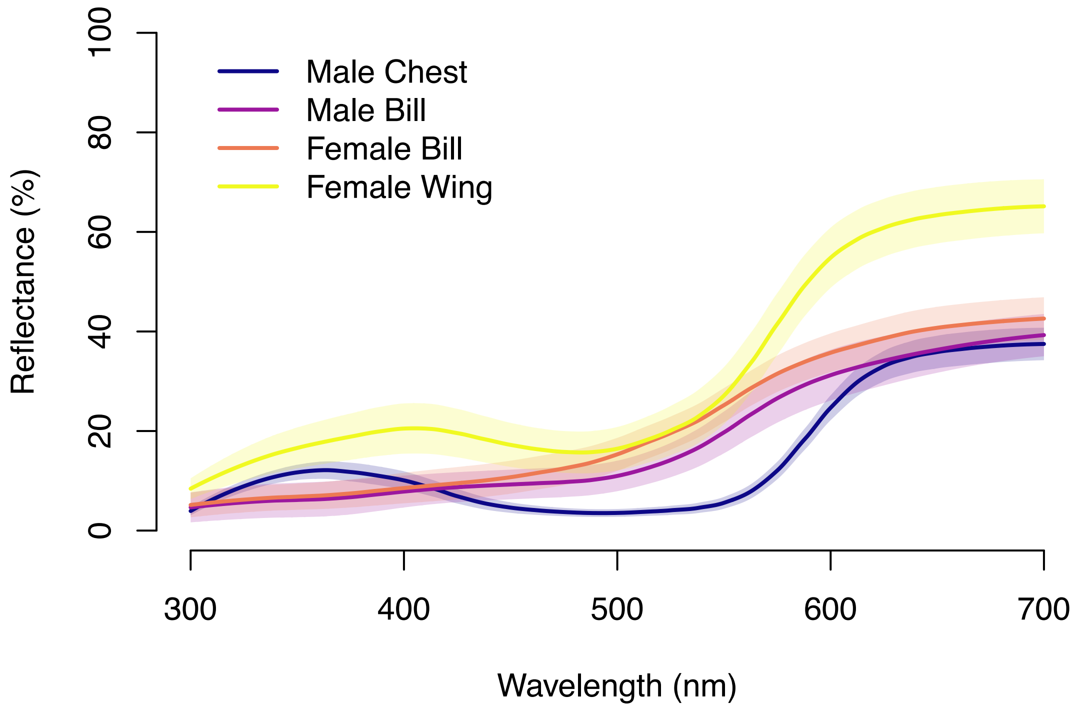


**Figure S1.** Examples of reflectance curves for the signals measured in the study. The dark line represents average reflectance and the shaded area represents one standard deviation. The male chest curve was generated from ten random males measured in the study. The male bill, female bill, and female wing curves were generated from ten measurements of museum skins. All signals reflect predominantly long wavelengths and have minimal ultraviolet (UV) reflectance.

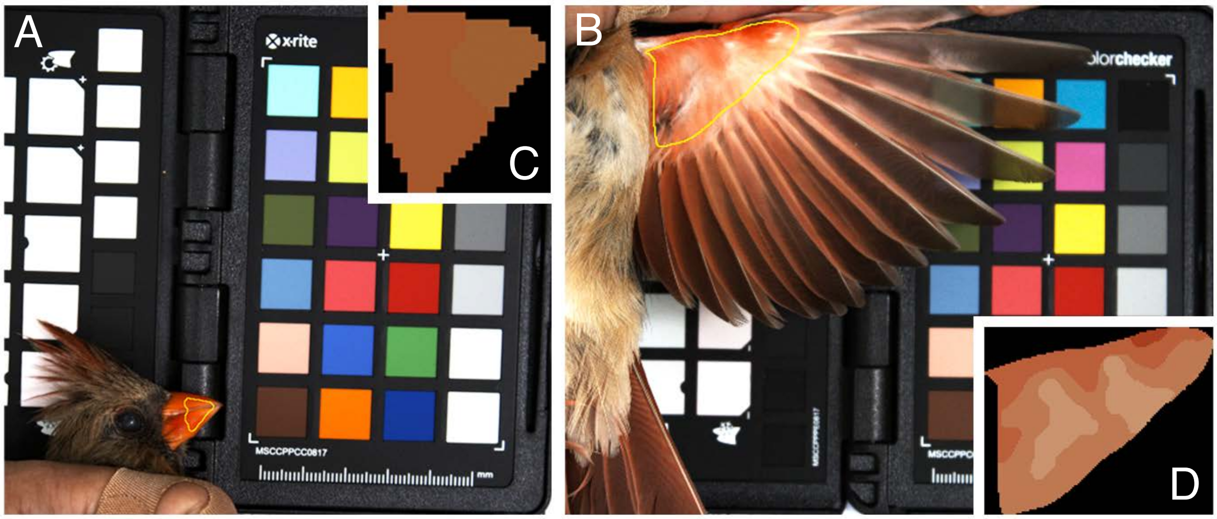


**Figure S2.** Examples representing the analysis of male and female bill (a) and female wing (b) photos. After selecting regions of interest, we used the QCPA framework to break the color patch up into discernable clusters (c, d) that could be measured for hue, saturation, and brightness.
